# NeoDBS: Open-Source Platform for Visualization and Analysis of Electrophysiological Recordings from Deep Brain Stimulation Systems

**DOI:** 10.64898/2026.03.27.714691

**Authors:** Larissa Rodrigues, Andreia Ferreira, Inês Pereira, Ricardo Moreira, Luis Jacinto

## Abstract

Optimization of deep brain stimulation (DBS) therapy for neurological and neuropsychiatric disorders depends on objective quantitative biomarkers that can guide stimulation parameter adjustments. With the recent introduction of new-generation DBS systems capable of simultaneously stimulating brain activity and recording local field potentials (LFP), there is increasing demand for platforms that enable efficient visualization and analysis of these signals for electrophysiological biomarkers identification. To address the limitations of currently available toolboxes that require advanced signal processing skills and rely on proprietary software, we present NeoDBS, an open-source Python platform designed for ingestion and advance signal visualization and processing of LFP signals from DBS systems through an easy-to-use graphical interface. NeoDBS is a user-centered platform that offers predefined analysis pipelines with the aim of facilitating electrophysiological biomarker investigation for DBS across different brain disorders. Custom analysis pipelines are also available for users to leverage the signal analysis tools to their research needs. Critical functionalities for longitudinal biomarker research are featured in NeoDBS, such as batch file processing and event-locked analysis for in-clinic and at-home recordings. This combination of accessibility, user-experience and advanced signal processing tools makes NeoDBS an environment that propels easy and fast electrophysiological biomarker research for DBS, across patients, sessions, and stimulation parameters.

## 1. Introduction

Deep brain stimulation (DBS) is an established neuromodulatory therapy for Parkinson’s Disease (PD) [1] and other movement disorders, such as essential tremor or dystonia [2], and an emerging treatment for severe and refractory cases of neuropsychiatric disorders, including obsessive-compulsive disorder (OCD) [3] and depression [4]. DBS is a highly individualized therapy involving the surgical implantation of electrodes that deliver electric pulses to modulated the activity of targeted brain regions, requiring careful adjustment of the stimulation parameters for optimal clinical outcome. Although DBS can significantly improve the quality of life of those afflicted by neurological and neuropsychiatric disorders, the achievement of optimal therapeutic effects without excessive side effects is reliant on a lengthy process in which patients must attend numerous follow-up appointments after surgery. In these appointments, stimulation parameters are manually adjusted by the physician, relying only on subjective self-reports from patients or clinical scoring scales [5]. The lack of robust biomarkers that correlate clinical efficacy to stimulation parameters or clinical states limits the clinical management of the disorders and the effectiveness of the treatment [6]. Identifying objective quantitative biomarkers that correlate with the patient’s clinical state, symptom evolution and therapy efficacy can ultimately guide optimization of stimulation parameters and advance of responsive stimulation [7].

Electrophysiological biomarkers derived from intracranial neural activity can serve this purpose, as neural activity can be actively modulated by symptoms, physiological states, behavioral events, and circadian rhythms. This is especially important for adaptive DBS (aDBS), an emerging closed-loop approach that can automatically titrate stimulation parameters directly from real-time neural activity biomarkers, correlated to disorder-related events or symptoms [8], [9], [10]. This configuration allows intermittent stimulation, leading not only to improved therapeutical results but also a reduction of side effects and battery consumption [11]. aDBS is made possible through the use of new generation DBS systems that are capable of simultaneous stimulation and recording of deep brain activity. At present, and since 2020, the only DBS systems approved for clinical use in the European Union (EU) and in the United States (USA), are the Medtronic Percept™ PC/RC neurostimulators with BrainSense™ technology, differing only on the type of battery integrated in the neurostimulator. These systems can record local field potentials (LFPs), which are extracellular electrophysiological potentials reflecting the coordinated activity of a large population of neurons, from DBS leads [12]. LFPs have been used for decades for the investigation of brain disorders in preclinical animal models, but also in human studies mostly from interoperative electrophysiological recordings, and have been correlated with an array of cognitive and motor functions but also with pathological states across different brain disorders [13], [14], [15]. Recently, aDBS has been approved for medical use in Parkinson’s disease (PD), where features from beta band oscillations extracted from LFP signals from the subthalamic nucleus (STN) serve to automatically adjust stimulation amplitude and improve motor function [8], [11].

The identification of reliable electrophysiological biomarkers from LFPs recorded from DBS systems is a critical step for the wider adoption of aDBS for other neurological and neuropsychiatric disorders beyond PD. However, extracting reliable quantitative metrics from LFPs typically requires significant knowledge in signal processing and programming. This presents a substantial challenge to clinicians and researchers lacking expertise in these domains, limiting their ability to transform raw intracortical signals into quantifiable metrics that reflect neural processes. Although previous efforts have been made in the development of toolboxes for visualization and analysis of LFPs recordings from DBS systems, such as Percept Toolbox [16], Perceive Toolbox [17], DBScope [18], and BRAVO [19], these were limited by the reduced number of functionalities, paid development environments, or absence of user-friendly interfaces and analysis pipelines.

Here, we present NeoDBS, a free open-source Python-based toolbox for offline data visualization, processing, and feature extraction from Percept™ PC/RC + BrainSense™ recordings. The novel toolbox addresses limitations of previously developed tools and includes batch processing of multiple recording sessions, event-locked analysis for chronic and in-clinic data, and automatic feature extraction, in an intuitive browser-like interface with predefined pipelines for users lacking signal processing skills (Figure 1).

**Figure 1.**
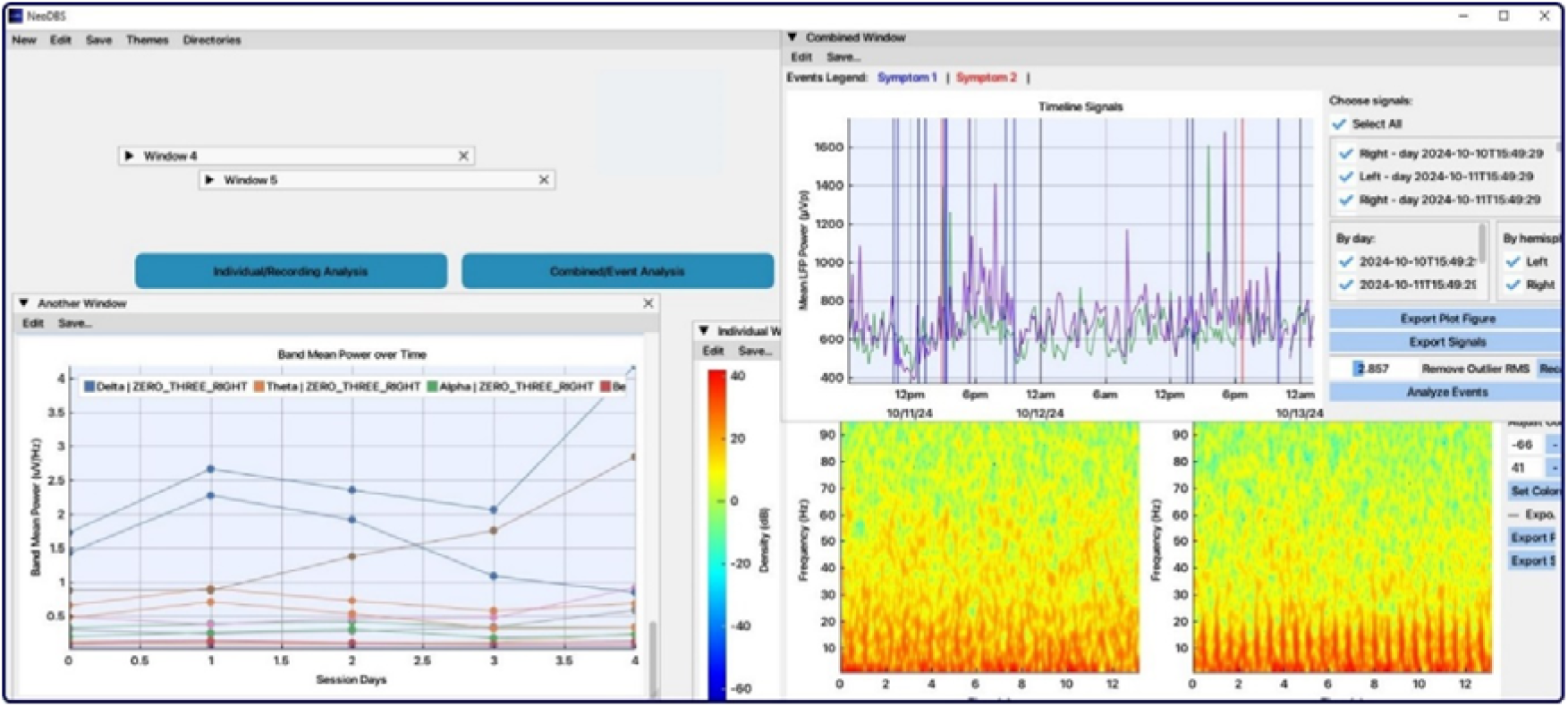
NeoDBS graphical user interface (GUI) showcase. With a browser-like design, NeoDBS enables multiple workspaces (windows) destined for the analysis of DBS recordings, that run independently and simultaneously. This design is meant to propel longitudinal and comparative analysis across patients and/or recording sessions.

## 2. Methods and Results

NeoDBS was developed in Visual Studio Code (Windows x64), with Python v3.11.9. The source code and documentation for NeoDBS, including standalone installers for Windows, Mac, and Linux are publicly hosted on a GitHub repository (https://github.com/neuroephys-lab/NeoDBS). Full documentation includes installation steps and tutorials to help users explore NeoDBS’s functionalities, as well as descriptions of all signal processing functions. Besides custom-written functions, NeoDBS uses functions from the following Python libraries: Dear PyGui [20] (for graphical interface); SciPy [21] and NumPy [22] (for signal processing), and MatplotLib [23] (for Figure export). DBS recordings from Medtronic Percept™ PC/RC neurostimulators + BrainSense™ technology were used to develop and validate NeoDBS. Signals were acquired from patients with refractory obsessive-compulsive disorder followed at University Hospital Center of São João (Porto, Portugal).

### 2.1. Types of Recordings

Medtronic Percept™ + BrainSense™ system is the only DBS system approved for clinical use in the EU and USA for simultaneous stimulation and recording of neural activity [12], [24]. Thus, NeoDBS was designed, at this stage, to ingest, process, visualize, and analyze recordings from this system. Medtronic BrainSense™ technology features different recording modes for different purposes and configurations. NeoDBS treats each recording mode independently so that predefined analysis pipelines tailored to each mode can be presented. Recordings modes can be divided into two main categories, according to the context in which they were acquired: in-clinic recordings or at-home recordings [12], [25].

In-clinic recordings include the recording modes BrainSense™ *Streaming, Indefinite Streaming, Survey*, and *Setup*, which are acquired in an in-clinic context, typically in the presence of the physician. They can be used during patient appointments to analyze the patient’s neural activity with or without active stimulation therapy, but also to assess electrode integrity, configure stimulation and recording leads, and identify artifacts. The streaming modes record neural activity in real-time at 250 S/s, for as long as desired, and can be used for clinical research purposes while the patient performs protocol tasks or responds to different stimulation parameters, allowing the study of real-time modulation of the neural signals [15].

At home-recordings include BrainSense™ *Timeline* and *Events* modes and are available when chronic sensing has been enabled by the physician to allow long-term (24/7) monitoring of the patient’s neural activity outside of clinic visits. BrainSense™ *Timeline* signals contain the averaged power of a predefined 5 Hz band, calculated every 10 minutes. In the *Events* mode the patient can record specific events, such as symptoms or medication intake, with the use of Medtronic’s patient programming app. These events activate a snapshot of brain activity in the form of a power spectrum derived from 30s of neural activity recorded at a sampling rate of 250 S/s from the event onset, triggered by the patient through the patient’s programming app. Signals from all recording modes can be visualized and/or analyzed with NeoDBS (see section 2.2). Table 1 summarizes NeoDBS’s signal processing and analysis features for each recording mode.

**Table 1:** Summary of recording modes, data types from BrainSens™ technology, and respective nomenclature and available functionalities to process these recording modes in NeoDBS.

| Categories | System Name | NeoDBS Name | NeoDBS Functionalities |
| --- | --- | --- | --- |
| **In-Clinic Recordings** | BrainSense™ *Streaming* | *Streaming* | Segmentation and filtering; Cardiac artifact removal; Event annotation; Event-locked segmentation; Power spectral density (PSD) and coherence estimation; Spectrogram, Wavelet and Coherogram; Phase-amplitude coupling (PAC); Cross-correlation; Automatic feature extraction. |
|  | BrainSense™ *Indefinite Streaming* | *Indefinite* |  |
|  | BrainSense™ *Survey* | *Survey* |  |
|  | BrainSense™ *Setup* | *Setup* |  |
| **At-Home Recordings** | BrainSense™ *Timeline* | *Timeline* | Circadian rhythm display; Event-locked segmentation |
|  | BrainSense™ *Events* | *Events* | Automatic feature extraction |

**Table 1:**
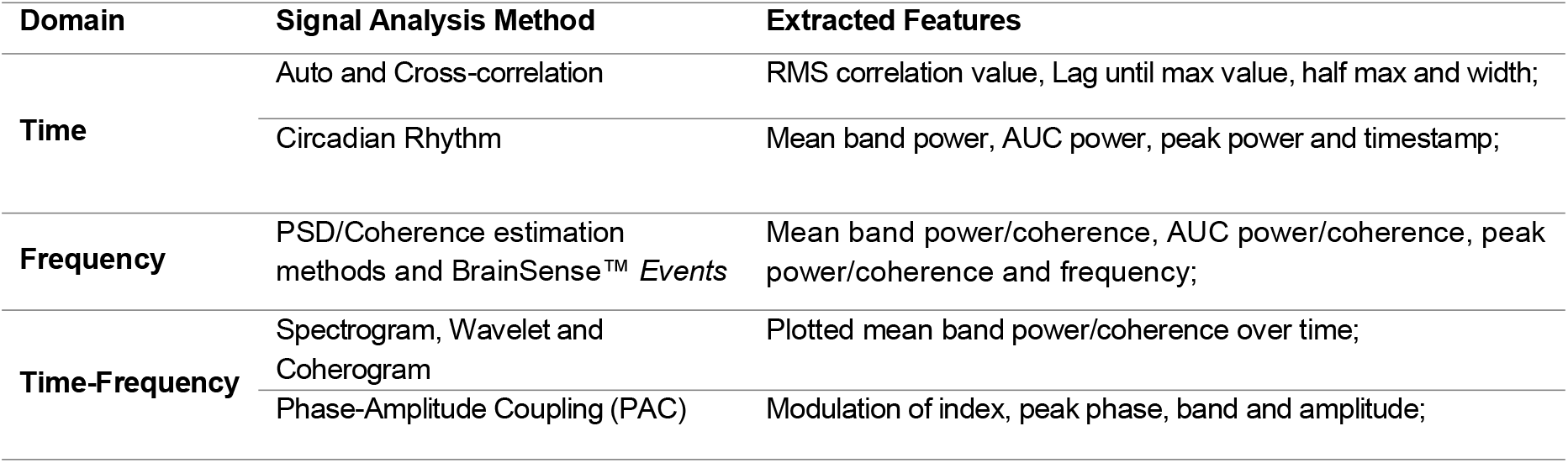
Extracted features per signal analysis method, in NeoDBS.

| Domain | Signal Analysis Method | Extracted Features |
| --- | --- | --- |
| Time | Auto and Cross-correlation | RMS correlation value, Lag until max value, half max and width; |
|  | Circadian Rhythm | Mean band power, AUC power, peak power and timestamp; |
| Frequency | PSD/Coherence estimation methods and BrainSense™ *Events* | Mean band power/coherence, AUC power/coherence, peak power/coherence and frequency; |
| Time-Frequency | Spectrogram, Wavelet and Coherogram | Plotted mean band power/coherence over time; |
|  | Phase-Amplitude Coupling (PAC) | Modulation of index, peak phase, band and amplitude; |

### 2.2. Graphical interface and workflow

NeoDBS’ graphical user interface (GUI) was designed as a browser-like platform, in which multiple concurrent analysis can be run simultaneously in independent windows (Figure 1). In practice, each window works as an independent workspace for signal visualization and analysis, allowing users to explore different signal processing and quantification routes and compare results, e.g. in windows side-by-side. The workflow inside a window follows a window (file) - tab (recording) - child tab (analysis) hierarchy (Figure 2). After opening a specific file, or set of files, in a new window (Figure 2A), the user can access each type of recording modes available within the files (see section 2.1) as independent tabs (Figure 2B). Inside each tab, plots of the respective signals from all available electrodes are shown, as well as all available signal processing and analysis functions for that type of recording (Figure 2B). Signal segmentation and filtering is possible at this stage, before the application of further analysis (Figure 2C). When a signal analysis method is chosen within each signal tab, a child tab is created with the respective representation or output of the selected method (Figure 2D). Metrics are automatically extracted and organized into tables (Figure 2E), and are easily exported (see section 2.9). The user can then create as many child tabs as needed for parallel analysis of the same signal.

**Figure 2.**
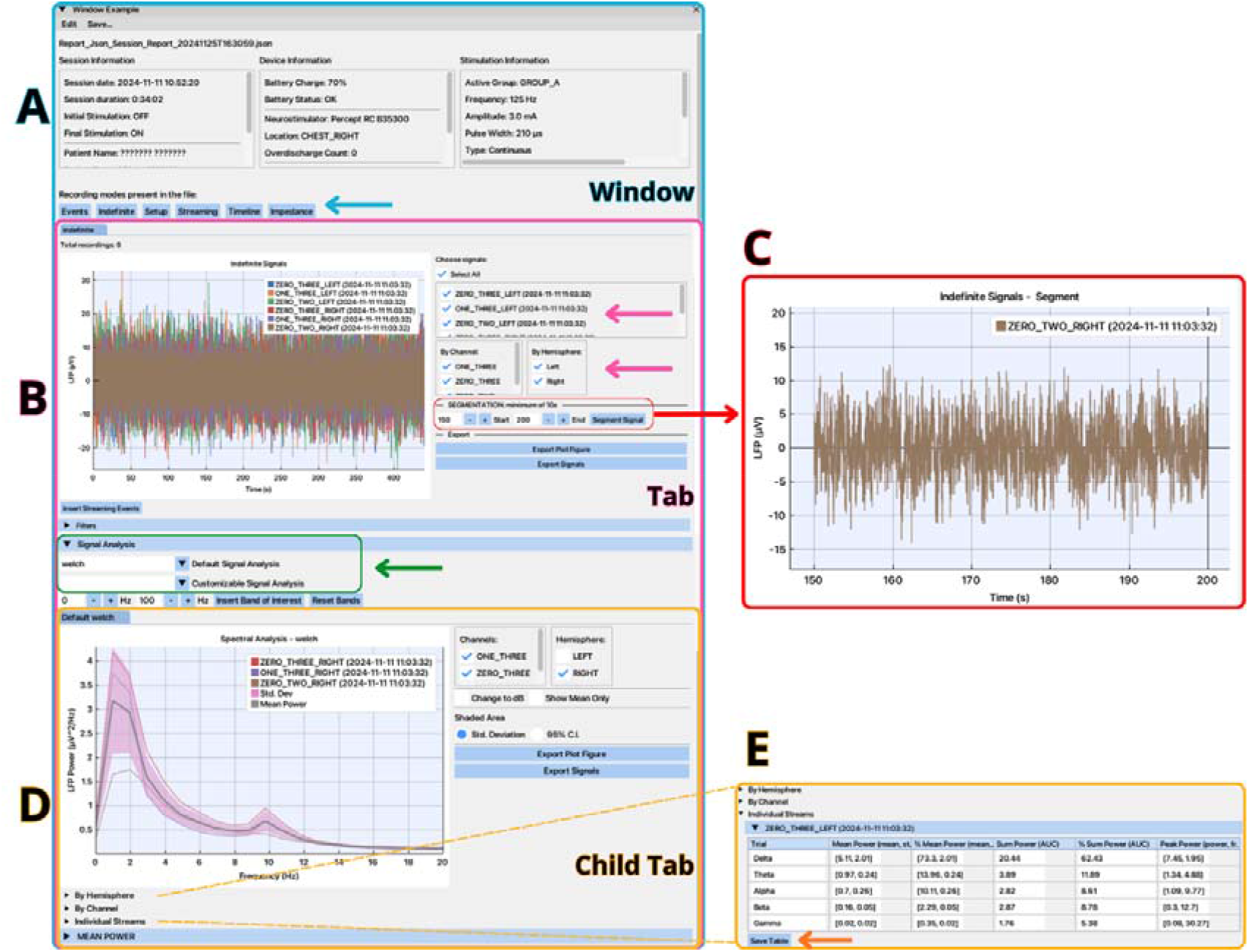
NeoDBS window workflow example. A) In the base window, after selecting which file to open, NeoDBS presents information regarding the recording session(s) including patient, device, and stimulation parameters, and displays the recordings modes containing signals within the file (blue arrow). B) After the user presses the recording mode of interest, a tab will open with a time-domain plot of the signals, for all available electrodes, and present which functionalities are available for processing and analyzing those signals. As an example, a plot of 6 *Indefinite Streaming* signals from 6 electrode pairs (3 from each hemisphere) are shown. The user can toggle which signals from each electrode are visible in the plot (pink arrows) and C) perform segmentation of the selected signals, for e.g., selecting only 50 seconds from one 450 seconds recording from one electrode pair, as shown. D) By choosing a signal analysis function from a drop-down menu (green arrow), a child tab is created with the output of the method. E.g. a power spectral density (PSD) estimation of the right hemisphere signals present in B, calculated by the Welch method, showing the individual PSD estimate for each electrode pair as well as the mean PSD and respective standard deviation for the three electrode pairs. E) Features from the PSD, including absolute and normalized mean power, area under the curve, and peak power and its frequency, are automatically extracted and presented in a table. All signals, figures, and tables from any tab can be exported to files and saved on the hard drive (orange arrow).

### 2.3. Types of windows

NeoDBS permits the analysis of single individual files, typically from single in-clinic sessions or containing at-home recordings from a specific period, or batch processing of multiple files from multiple in-clinic sessions or multiple at-home recordings. The capability to extract metrics from LFP signals across time and/or patients is fundamental to increase statistical confidence on any observed clinically-relevant electrophysiological biomarkers but also to assess how these biomarkers are modulated over time and by the stimulation [9], [26]. Thus, there are two types of main analysis windows in NeoDBS: the Individual window and the Combined window. As the name reflects, the Individual window allows the ingestion of a single individual file, focusing on the evolution of signals and events inside the same recording session. The Combined window allows the selection of multiple files, focusing therefore on the evolution of the signals across different recording sessions, patients, and protocols or the extraction of mean metrics across all selected sessions. The available functionalities within NeoDBS for each type of window are summarized in Table 2.

**Table 2:**
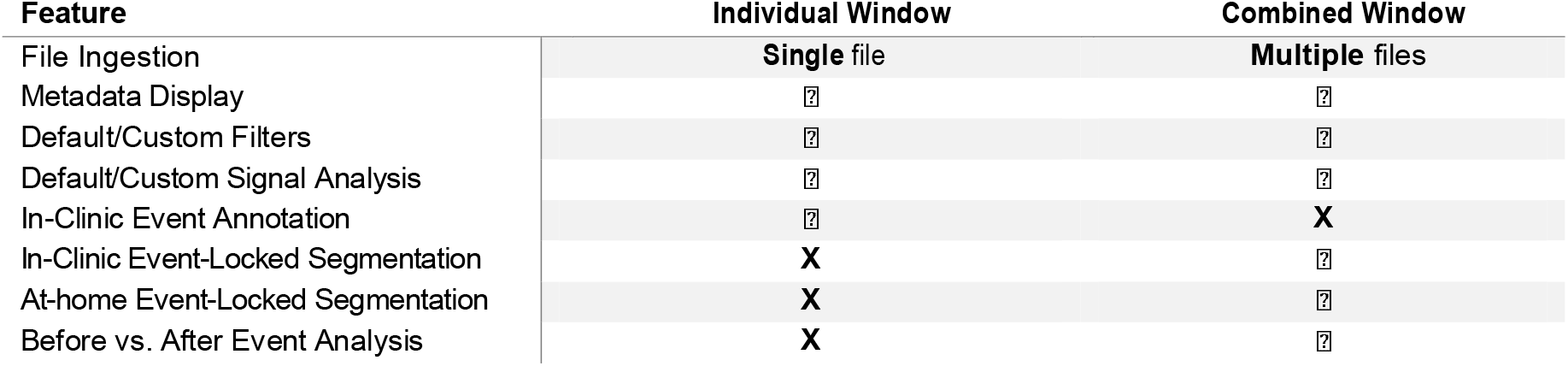
Summary of available functionalities per workspace (window).

**Table 2:** Comparison of functionalities between NeoDBS and available open-source platforms for LFP-DBS visualization and analysis.

| Platform | Interface | Data Visualization and Analysis | Event-Locked Analysis | Feature Extraction | Batch Analysis |
| --- | --- | --- | --- | --- | --- |
| **NeoDBS** | ☑ | All recording modes | In-clinic and At-home data | Automatic feature extraction, exportation available on GUI | ☑ |
| **Perceive Toolbox** | ☑ | In-clinic *Streaming* data | **X** | ☑ | **X** |
| **Percept Toolbox** | **X** | All recording modes | At-home data | **X** | **X** |
| **DBScope** | ☑ | At-home data, *Streaming* and impedance values | At-home data | **X** | Only ingestion |
| **BRAVO** | ☑ | Available for in-clinic and at-home data | At-home data | Automatic feature extraction | Only ingestion |

### 2.4. Analysis pipelines

One of NeoDBS’ main objectives is to allow users without previous knowledge of signal processing methods to be able to explore recordings from DBS systems with the tools integrated in the platform. Therefore, all the filters and signal analysis methods (described in sections 2.5, 2.6, and 2.7) have default and custom pipelines. In the default pipelines the methods’ parameters are predefined with values from predefinitions of the libraries used, or hard coded with suitable values to the acquisition configuration. Custom pipelines allow users to fully define all the parameters in the user interface, only restricted by the selected function requirements. To guide user in the fine-tuning of parameters, documentation for each method can be accessed inside the platform and presents the description of all tunable parameters within a function/method.

### 2.5. Data segmentation and filtering

In-clinic and at-home recordings can capture capture modulations of the neural activity relating to relevant clinical events, such as behavioral or symptom changes. Segmentation of epochs of interest is thus critical for analysis of these modulations for event-related biomarker identification. For data segmentation, NeoDBS includes both a manual segmentation function, where the user can isolate selected segments from signals with a minimum of 10 seconds of duration, and an automatic segmentation function which uses timestamps to automatically create event-centered epochs with a user-defined duration (see section 2.6 for event-locked analysis methods).

Following segmentation of segments of interest, filtering LFP signals is a standard preprocessing step to either remove sources of noise or to isolate oscillatory brain activity within frequency bands with physiological relevance, such: delta (0.5-4 Hz), theta (4-8 Hz), alpha (8-12 Hz), beta (12-30 Hz), and gamma (30-100 Hz) bands [11], [27], [28]. Filters available in NeoDBS include several infinite impulse response filters (IIR) such as Butterworth, Chebyshev I and II, ellipsoid, Bessel, notch and peak, with user-defined cut-off frequencies.

LFPs collected from DBS systems can also be contaminated by noise from different sources including stimulation, motion, and cardiac artifacts [25], and present missing data due to temporary communication failure or internal hardware data processing failure [16]. To reliably extract meaningful task or clinical-related metrics from intracranial LFP signals it is important to investigate for the presence of these artifacts and apply signal processing techniques that can isolate the signal of interest without distortion. To remove added sinusoidal noise, such as 50/60 Hz line noise, the user can simply apply any of the filters described above, adjusting the cut-off frequencies. Cardiac artifacts are also common in LFPs recorded from DBS systems due to neurostimulator implantation close to the cardiac dipole, being more prevalent in recording configurations where the stimulation is enabled [29]. The cardiac artifact rejection method used in NeoDBS was adapted from Bravo [19], and is applied when an electrocardiogram (ECG) template is identified, through peak method and subtracted from the raw signal, posteriorly filtered with Butterworth bandpass.

Timeline recordings also regularly present outlier values that can be identified as “overvoltage events” due to device’s saturation of the analog-to-digital converter [30]. These artifacts, which have a much higher magnitude than the signal’s mean, disrupt the visualization and interpretation of *Timeline* data [25]. In NeoDBS, the user can filter outlier values using a custom outlier removal tool based on a user-defined threshold above the root mean square (RMS) of the recording. Additionally, overvoltage events can ultimately lead to missing samples [25], but NeoDBS automatically inspects *Timeline* signals for missing samples within the recording period, alerting the user when they occur and marking them on the signal plot. A mean interpolation function is then available if the user desires to interpolate missing samples.

### 2.6. Event-locked analysis

Event-locked analysis can be used to identify clinically relevant metrics that are correlated with specific physiological and behavioral events. NeoDBS includes event-locked analysis functions for both in-clinic and at-home data. Two types of events can be analyzed in NeoDBS: clinician-defined in-clinic events and patient-marked at-home events.

The in-clinic recordings modes in Percept™ PC/RC + BrainSense™ do not allow for direct system annotation of events, either by the patient, the clinician, or from external device triggers, which can limit the analysis of neural activity related to clinically relevant events. As an alternative, NeoDBS allows users to upload a text file with timestamps (in UTC format) of events that occurred during the recording session, coded by event type, and displays them on top of the signal plot.

An example of in-clinic event annotation is represented in Figure 3A, with an obsession and a compulsion event marked by the clinician in UTC format during a clinical evaluation of an OCD patient. clinician-defined event annotation is possible for both single recording sessions and multiple recordings session in batch processing (Figure 3A-B). The latter allows, for example, the analysis of mean responses to a specific event across different recording sessions (Figure 3C).

**Figure 3.**
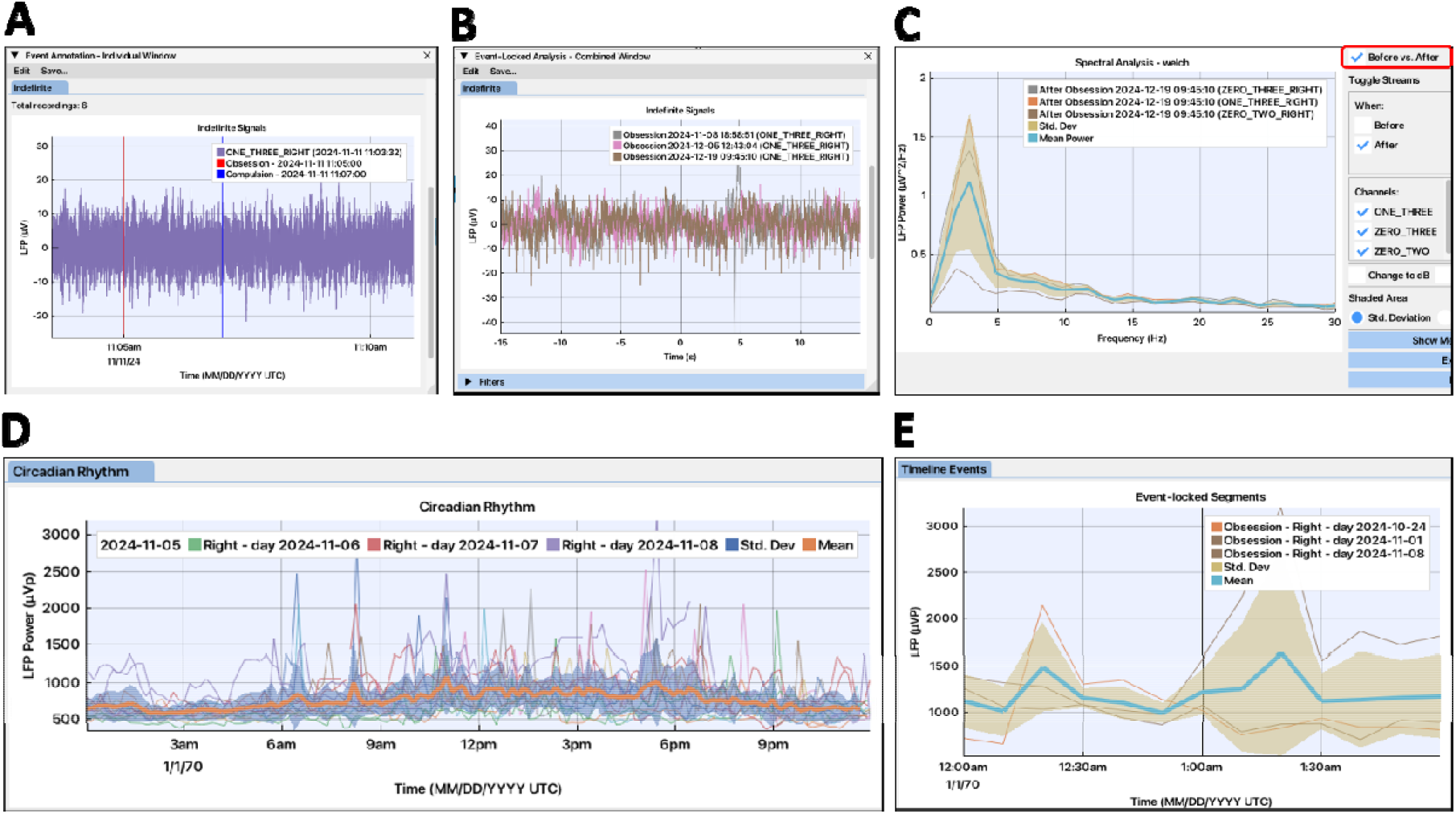
NeoDBS event-locked analysis and circadian rhythm. A) Time-domain signal from one electrode pair with clinician-defined event annotation in Individual window. Vertical lines represent the timestamps of the events, including an obsession event (red) and a compulsion event (blue) in an in-clinic recoding from an OCD patient; B) Event-locked segmentation of multiple in-clinic recordings, in Combined window, showing signals recorded from the same electrode pair in three different sessions and centered at an obsession event; C) Mean power spectral density (PSD) estimate calculated from signals from one electrode during three obsession events previously segmented for event-locked analysis; D) Circadian rhythm with mean power and respective standard deviation over a 24 hour period extracted from multiple days from BrainSense™ *Timeline* recording; E) Event-locked segmentation using BrainSense™ *Events* timestamps to calculate the mean power of three BrainSense™ *Timeline* recordings in a 2-hour epoch centered at an obsession event.

For at-home recordings, NeoDBS allows the visualization of the mean circadian rhythm extracted from the BrainSense™ *Timeline* by averaging power estimates across 24 hours from multiple days (Figure 3D). Event-locked analysis of patient-marked events from at-home recordings is also possible by the creation of mean power epochs centered at the chosen events from BrainSense™ Events timestamps (Figure 3E).

### 2.7. Time and spectral analysis tools

Intracranial neural signals are highly non-stationary and characterized by signal properties that change over time. The frequency-domain component of LFPs recorded from DBS systems can be investigated through power spectral density (PSD) and coherence estimation tools. PSD estimates the distribution of energy across the frequencies of a signal, providing an interpretable measure of neural oscillatory activity across recordings or events. For example, power variations in beta band frequency in the SNT of PD patients has been established as an electrophysiological biomarker correlated to motor symptoms [2], [8]. NeoDBS offers PSD estimation using the periodogram, Welch’s, and multitaper methods (Figure 4A), with each method offering different trade-offs between frequency resolution estimate variability [29], [31], [32]. Because PSD estimates do not account for time-based modulation of signals across time, NeoDBS allows users to analyze time-frequency modulations of the neural signals by employing spectrograms (Figure 4B) or wavelet (Figure 4C) methods. These signal representations are important to examine how the oscillatory activity of the different frequency bands is modulated across time, for example during adjustment of stimulation parameters [26] or symptom provocation protocols [33]. Additionally, in NeoDBS, computed band power over time can be extracted from the spectrograms and plotted in isolation, facilitating the quantitative analysis of how oscillatory brain activity for each band fluctuates between tasks and events.

**Figure 4.**
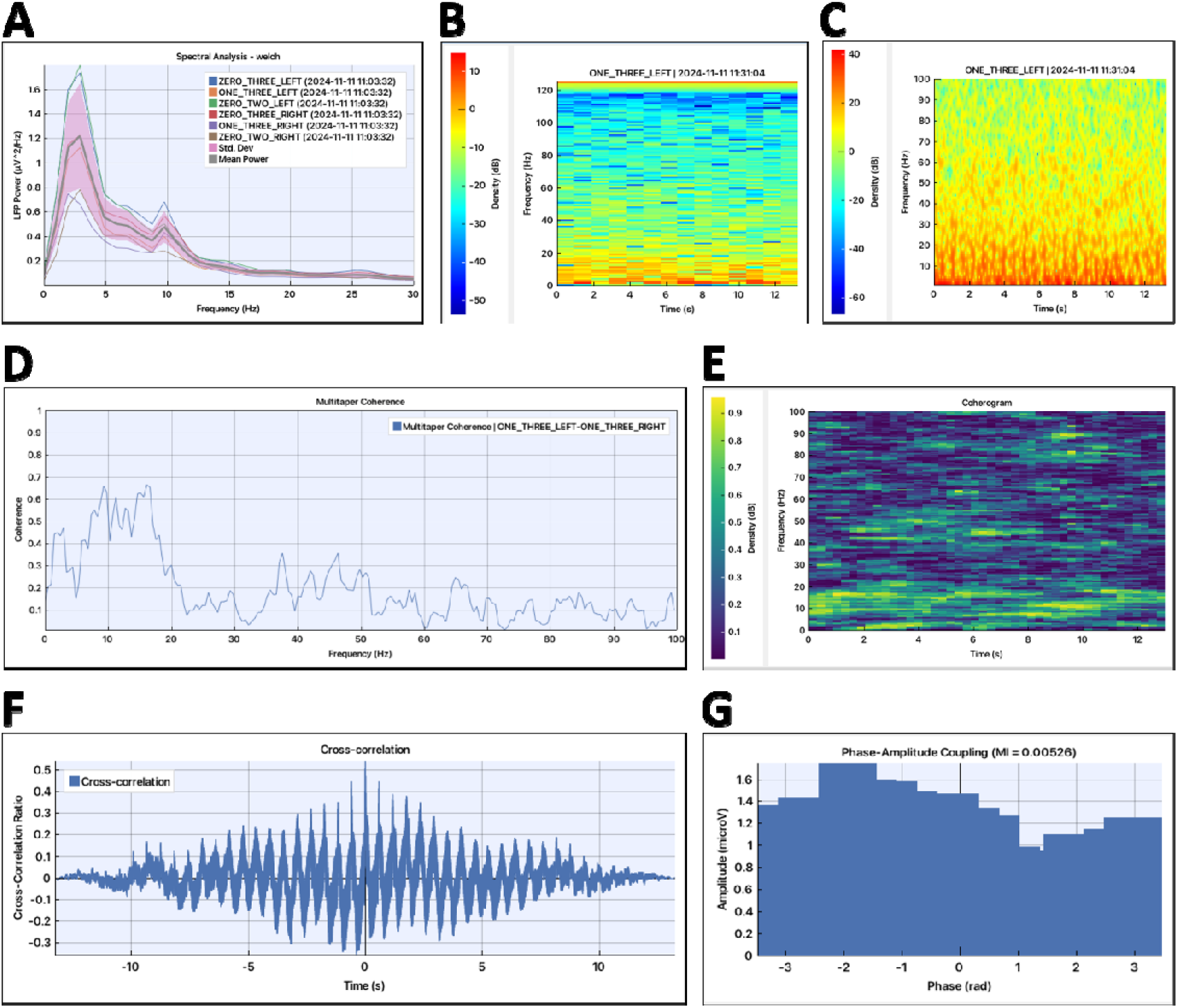
Main NeoDBS functionalities available for the analysis of in-clinic recordings. A) PSD estimation with multitaper method for six recordings from six electrode pairs from one in-clinic *Streaming* recording session; B) Spectrogram and C) Wavelet spectrogram (or scalogram) from one electrode pair from one in-clinic *Streaming* recording; D) Coherence magnitude computed between two *Streaming* electrode pairs, from the same recording session; E) Coherogram computed between two *Streaming* electrode pairs, from the same recording session; F) Cross-Correlation between two *Streaming* electrode pairs, from the same recording session; G) Phase-amplitude coupling for delta (phase) and alpha (amplitude), between two *Streaming* electrode pairs.

Coherence analysis allows quantifying correlations between two signals in the frequency-domain, by calculating the strength of the linear relationship between the two signals at each frequency. This metric can be useful to assess functional connectivity and coordinated oscillatory activity, between hemispheres or different brain regions, both during resting-state and in response to stimulation or clinically-relevant events [34], [35], [36]. NeoDBS includes coherence computation with Welch and multitaper PSD estimation (Figure 4D). Coherogram estimation, which calculates how coherence is modulated over time across multiple frequencies, is also available (Figure 4E).

Time-dependent modulations of LFP signals can be examined in NeoDBS using autocorrelation and cross-correlation calculations. Autocorrelation quantifies the similarity of a signal with a time shifted version of itself, allowing identification of repeating temporal patterns or rhythmic structure within a single recording. In contrast, cross-correlation measures the similarity between two different signals as a function of temporal lag, enabling detection of directionality and time-dependent synchronization of neural activity between different recording sites (Figure 4F). These approaches can be useful to uncover temporal synchrony or patterns within brain regions, or study functional connectivity and dependency across different brain regions, respectively [37], [38].

Phase-amplitude coupling (PAC) correlates the phase of a specific frequency range with the modulation of amplitude of the same or other frequency ranges. PAC analysis has been previously used to show, for example, that STN DBS can alter beta phase-locking between the basal ganglia and the motor cortex, and that these changes correlate with clinical motor symptoms in PD [39]. PAC estimation is available in NeoDBS for signals from the same electrode pair or between signals from different electrode pairs (Figure 4G).

### 2.8. Feature extraction

To identify and validate neural biomarkers, it is essential to extract quantitative features from signal transformations and analyses, enabling objective characterization of neural modulations across events, tasks, or days. When any signal analysis method, described in section 2.7, is applied, NeoDBS automatically extracts a set of quantitative features, according to the type of analysis employed. The extracted features are shown organized in a table divided by groups (hemisphere, channel and/or event). For frequency-dependent analysis, the features of the delta, theta, alpha, beta and gamma bands are extracted by default. The user can also add custom frequency bands of interest for feature extraction, with a minimum of 2 Hz bandwidth. All features automatically extracted by NeoDBS for each signal analysis method, are summarized in Table 3.

### 2.9. Customization and data export

During NeoDBS usage, users can customize the names of windows, patient-marked events, recording channels, and select different fonts and colors. To facilitate the use of data generated in NeoDBS in other data analysis or image processing software, it is possible to export all data, including plots, signals, and tables in an organized fashion. Computed outputs can be exported as figures (for plots) or as comma separated value (csv) files (for processed signals and extracted metrics) for posterior offline analysis. Data export can be performed for each individual item, for each window, or for all existing items at once.

## 3. Discussion

Optimizing deep brain stimulation therapy requires uncovering the neural mechanisms that respond to stimulation parameters or symptom evolution. The analysis of ongoing neural activity recorded from new generation DBS systems that allow for simultaneous stimulation and recording of brain activity are an attractive path to analyze how neural signals correlate with symptoms, circadian rhythms, and stimulation parameters. This is particularly critical for the advent of aDBS, in which the same neural signals can guide automatic adaptive adjustments of the stimulation parameters, improving treatment efficacy, and reducing the burden on clinicians and healthcare providers [33]. In this work, we presented a novel open-source platform that aims to facilitate the analysis of complex neural signals recorded from new generation DBS systems, and advance the identification of relevant electrophysiological biomarkers for disorders that benefit from DBS therapy.

For clinicians and researchers, sharing knowledge and advancing methodologies is essential for analysis standardization, biomarker validation, and therapy/research optimization. Free open-source platforms are instrumental in facilitating dissemination and collaboration by enabling the exchange of standardized quantitative information between research laboratories. Previous analysis platforms for signals from DBS systems were released as open-source tools [16], [17], [18], [19] but most have relied on MATLAB, a proprietary data processing environment, which can create cost barriers to wide adoption. NeoDBS was developed with Python, an alternative open environment that is both free of charge and well-suited for community development. One previous analysis platform, BRAVO, was also developed in Python, but it required setting up a local server or using a cloud-based server which could impose limits on the sharing of confidential clinical data [19]. A comparison of the general functionalities between NeoDBS and previously available open-source platforms to process and analyze BrainSense™ technology recordings is present in Table 4.

Another major limitation preventing widespread use of these platforms is the lack of a simple graphical user interface with accessible default functions, demanding that users have previous experience in signal processing and/or coding to implement all available functions. NeoDBS allows unexperienced users to implement all signal processing and data analysis methods available with default parameters. This means that users with no previous knowledge of signal processing methods are still able to visualize, analyze, and extract metrics from the LFP recordings. Advanced users can still take advantage of their signal processing skills by defining custom parameters for all functions. The design of the graphical interface of NeoDBS also considered user experience, and created a windows-based approach, that allows that multiple analysis windows and signals are opened simultaneously, which is familiar to most computer users with a standard operating system.

NeoDBS also implements novel batch processing in LFP analysis by allowing users to employ all analysis methods to simultaneously process several recording sessions in one single workspace and extract combined metrics. This functionality offers standardization of analysis across recording sessions, patients and tasks, while also allowing robust metrics extraction from longitudinal modulations of neural signatures. The combination of batch processing with event-lock analysis in NeoDBS also facilitates the identification and validation of electrophysiological biomarkers correlated with specific symptoms across patients, recording sessions, and protocol tasks.

One limitation of NeoDBS is that it can only handle files from the Medtronic Percept™ PC/RC + BrainSense™ systems. Although this is the only system currently approved for medical use in the EU and USA that can combine stimulation and recording capabilities, other devices from other manufacturers are set to emerge in coming years, especially as aDBS is further approved for other disorders beyond Parkinson’s Disease [11], [33], [40]. Since most signal processing and data analysis functions available in NeoDBS can be applicable to any neural signal from any system, file loaders for each specific system can be added to the platform without changing its main workflow. Another limitation of NeoDBS is the lack of a dedicated local database, with data managed in memory by the platform. While data integrity and persistence are ensured, this design can lead to performance degradation when computationally intensive tools are used, as memory and processing resources are more heavily taxed. It is recommended that computationally intensive methods such as time-frequency representations are used in smaller segments (under 10 minutes).

NeoDBS is under continuous development, including interface optimization and the addition of new tools which may expand to handling other types of signals, such as electroencephalogram and intraoperatory recordings, that can be correlated with DBS stimulation and recorded neural signals. New versions will remain open-source, empowering developers to modify the platform to their own needs and requirements. As active users increase, we expect that NeoDBS can contribute to the identification of electrophysiological biomarkers relevant to different neurological and neuropsychiatric disorders, driving DBS optimization and future closed-loop configurations.

## Ethics statement

Ethical approval for use of electrophysiological recordings from patients implanted with DBS systems was obtained from the ethics committee at University Hospital Center of São João (approval number 93/24). All participants provided their written informed consent prior to participating. The study conformed to the standards set by the Portuguese Law 21/2014 regulating clinical research, the EU’s General Data Protection Regulation (GDPR) 2016/679, and the Declaration of Helsinki.

## Acknowledgments

The authors would like to thank: the patients that contributed to this study; all units of University Hospital Center of São João where the patients were followed; and all members of the Neurophysiology and Neuroengineering lab for helpful discussions and for testing the multiple versions of NeoDBS. This project was funded by the Portuguese Foundation for Science and Technology (FCT) project 2024.17843.PEX.

## Author Contributions

LR and LJ conceptualized the platform. LR designed the graphical interface, programmed the main code, and implemented signal processing and data analysis functions. AF and IP contributed with signal processing and plotting functions. RM followed patients and performed electrophysiological recordings. LR analyzed patient data. LJ supervised the work. LR wrote the first draft of the manuscript. RM and LJ revised the manuscript. All authors read and approved the final version of the manuscript.∼

